## Supplemental Methods and Figures for "ZETA: a parameter-free statistical test for neuronal responsiveness"

### Supplementary methods

#### A proof of time-invariance

We have described some properties of the ZETA-test in the main method section, but we have not yet explained what the function is of the mean-subtraction of $\delta$ in eq. 14. This step plays a critical role in ensuring that the ZETA-test is time-invariant: i.e., that the latency of a neuronal response with respect to the stimulus onset does not affect the statistical significance of the ZETA-test.

We can see that this is the case if we have made a specific choice for the trial onsets, consisting of consecutive intervals of $\tau$, and made a set of $n$ spike times $v_1$ to $v_n$ in the interval $[0, \tau]$. First we rewrite equations 1 – 4 as:

$$\delta_i = \frac{i}{n} - \frac{v_i}{\tau} \qquad \text{S1}$$

Recall that $d_i = \delta_i - 1/n \, \Sigma_i \, \delta_i$. (eq. 6-7). Now, consider a shift of the trial onset times by $\Delta$ and let $v_k$ be the highest spike time smaller than $\Delta$. This results in a new set of $n$ spike times $v_i'$:

$$v_i' = v_{i+k} - \Delta \quad \text{for } 1 \leq i \leq n-k \qquad \text{S2}$$

$$v_i' = v_{i+k-n} - \Delta + \tau \quad \text{for } n-k+1 \leq i \leq n \qquad \text{S3}$$

Note that S3 implies circular time with the recording wrapping back to the beginning at the end of all trials, which we assume here to keep $n$ constant. If we define $\delta_i'$, analogous to $\delta_i$, and use equations S2-3 we find:

$$\delta_i' = \delta_{i+k} - \frac{k}{n} + \frac{\Delta}{\tau} \quad \text{for } 1 \leq i \leq n-k \qquad \text{S4}$$

$$\delta_i' = \frac{i}{n} - \frac{v_{i+k-n} - \Delta + \tau}{\tau} \qquad \text{S5}$$

$$= \delta_{i+k-n} - \frac{k}{n} + \frac{\Delta}{\tau} \qquad \text{for } n - k + 1 \le i \le n$$

And if we subtract its mean, the constants are removed, and we get:

$$d'_i = d_{i+k} \quad \text{for } 1 \le i \le n - k \qquad \text{S6}$$

$$d'_i = d_{i+k-n} \qquad \text{for } n - k + 1 \le i \le n \qquad \text{S7}$$

The set of $d'_i$ are thus identical to the set $d_i$ except for a reordering. The maximum of the set $|d_i|$ will therefore also be the maximum of the set $|d_i'|$. This means that it does not matter when the trial onsets are taken to compute ZETA. Note that this is not the case if $\delta_i$ is used instead of $d_i$.

To illustrate this difference, we will now derive closed-form solutions for the expectation and variance of $\delta_i$ and $d_i$ in the specific case of step-wise changing Poisson-distributed spiking rates. Note that this section serves only to illustrate the above derivation (Eq. S1 - S7) for a specific case, so the reader may choose to skip the rest of this section without missing out on any particularly important comments.

First, recall the base variables: we use $\boldsymbol{v}$ as a vector of spike times relative to stimulus onset, with the total onset-to-onset epoch duration defined as τ (see equation 1). We will set the neuron's firing probability to be homogeneous with Poisson rate $\lambda$. Since exponentially distributed inter-spike intervals generate Poisson-distributed spike counts, we use:

$$S \sim \text{Exp}\left(\frac{1}{\lambda}\right) \qquad \text{S8}$$

Therefore, spike time $w_i$ is:

$$w_i = w_{i-1} + S_i \qquad \text{S9}$$

In the limit of large $n$, the incremental deviation $\dot{D}_i$ from an exactly uniform spike-time distribution (i.e., $1/\lambda$) is therefore:

$$\dot{D}_i = \frac{1}{\lambda} - S_i \qquad \text{S10}$$

In reality, however, the rate $\hat{\lambda}$ is estimated from data that are limited by the observation window τ, number of trials $m$, and observed number of spikes $n = m\tau\hat{\lambda}$. First we collapse all spikes over trials, following Eq. 1, such that $v_n=\tau$. Then we normalize by τ to make $v_n=1$, which means that we now have:

$$\sigma \sim \text{Exp}\left(\frac{1}{m\tau\hat{\lambda}}\right) = \text{Exp}\left(\frac{1}{n}\right) \tag{S11}$$

Generating spike times:

$$v_i = v_{i-1} + \sigma_i \tag{S12}$$

With incremental deviations:

$$\dot{\delta}_i = \frac{1}{m\tau\hat{\lambda}} - \sigma_i \tag{S13}$$

We can confirm this is correct by rewriting the total deviation $\delta_i$ at spike $i$ as:

$$\begin{aligned} \delta_i &= \sum_{j=1}^{i} (\dot{\delta}_j) \\ &= \frac{i}{m\tau\hat{\lambda}} - v_i \\ &= \frac{i}{n} - v_i \end{aligned} \tag{S14}$$

To compute a closed-form for the variance of $d$, we need the first and second central moments of $\delta$ (i.e., the mean and variance). $\delta$ depends on $\delta$-dot, the variance of which is:

$$\begin{aligned} Var[\dot{\delta}] &= Var\left[\text{Exp}\left(\frac{1}{n}\right)\right] \\ &= \frac{1}{n^2} \end{aligned} \tag{S15}$$

In the case for large $n$, each $\delta$-dot is an exponential random variable. As $\delta_i$ is a sum over $\delta$-dots, we can approximate its pdf as an Erlang distribution with scale parameter $k$ equal to the spike number $i$:

$$\Delta_i = \sum_{j=1}^{i} (\dot{\delta}_j) \tag{S16}$$

$$\Delta_i \sim \text{Erlang}\left(k = i, \lambda = \frac{1}{n}\right)$$

However, we have so far ignored that $\delta_i$ is mean-zero and fixed at 0 at t=0 and t=τ. For a Wiener process W, such fixed points are known as a Brownian bridge (Mansuy & Yor, 2008), which is described by:

$$B(t) = W(t) - \frac{t}{\tau} W(\tau) \tag{S17}$$

However, since the underlying stochastic process in our case is not a standard normal, but Erlang-distributed, we cannot directly apply the above equation. Instead, we found that with sufficiently large *n*, the behavior of $\delta$ is described by a weighted difference of two time-symmetric series of Erlangs (see Figure 2-Supplement1B). One series grows as eq. 16, whereas its symmetric counterpart shrinks as $k$=$n$-$i$:

$$\delta_i = \frac{n-i}{2n} \text{Erlang}(i, \lambda) - \frac{i}{2n} \text{Erlang}(n-i, \lambda) \tag{S18}$$

Finally, the Erlang distribution is a specific case of the gamma distribution with integer scale parameters. The central moments of the difference between two gamma distributions are given by (Klar, 2015):

$$\mu = \frac{\alpha_1}{\beta_1} - \frac{\alpha_2}{\beta_2} \tag{S19}$$

$$\sigma^2 = \frac{\alpha_1}{\beta_1^2} + \frac{\alpha_2}{\beta_2^2} \tag{S20}$$

Where α=k=i, β=1/λ, and the subscripts indicate the two distributions. Filling in the above parameters and weighting variables, we therefore get:

$$E[\delta_i] = \frac{n-i}{2n} i\lambda - \frac{i}{2n}(n-i)\lambda = 0 \tag{S21}$$

$$\text{Var}[\delta_i] = \frac{n-i}{2n} i\lambda^2 + \frac{i}{2n}(n-i)\lambda^2 \tag{S22}$$

$$= \frac{ni\lambda^2 - i^2\lambda^2}{n}$$

Now, we need to compute the variance over $d_i$, which is defined as the mean-subtracted $\delta_i$.

To simplify the following derivation, we will assume that time is circular (as in S3) and that jittering does not strongly impact the number of spikes *n*, so we can we treat *n* as a fixed number of samples. As shown in Figure 2-Supplement1C, these assumptions allow accurate estimations. Now, we consider $v_k$, $1 \leq k \leq n$ taken from a uniform distribution on [0,1] and ordered such that $v_1 \leq v_2 \leq \cdots \leq v_n$. Then the probability distribution of $v_k$ is

$$p(v_k) = \binom{n}{k}\binom{k}{1}\int_0^{v_k} dv_1 \dots \int_0^{v_k} dv_{k-1}\int_{v_k}^1 dv_{k+1} \dots \int_{v_k}^1 dv_n \qquad \text{S23}$$

$$= \frac{n!\,k}{(n-k)!\,k!} v_k^{k-1}(1-v_k)^{n-k}$$

We find that the expectation of $v_k$ and $v_k^2$ are

$$E(v_k) = \int_0^1 dv_k p(v_k) v_k = \frac{n!\,k}{(n-k)!\,k!}\int_0^1 dv_k\, v_k^k (1-v_k)^{n-k} = \frac{k}{n+1} \qquad \text{S24}$$

$$E(v_k^2) = \int_0^1 dv_k p(v_k) v_k^2 = \frac{n!\,k}{(n-k)!\,k!}\int_0^1 dv_k\, v_k^{k+1}(1-v_k)^{n-k}$$

$$= \frac{k(k+1)}{(n+2)(n+1)} \qquad \text{S25}$$

Now, redefine

$$\delta_i = \frac{i}{n+1} - v_i \qquad \text{S26}$$

Then using Eq. S24 and Eq. S25, we find

$$E(\delta_i) = 0 \qquad \text{S27}$$

$$E(\delta_i^2) = \frac{i(n-i+1)}{(n+1)^2(n+2)} \qquad \text{S28}$$

We find that the variance of $\delta_i$ is parabolic with respect to i, with its minimum at $i = 1$ and $i = n$ and its maxium at the middle between those points. The maximum of all $\delta_i$ is thus also more likely to at the middle $\delta_i$ than at the extremes.

We define again

$$d_i = \delta_i - \bar{\delta} \qquad \text{S29}$$

with $\bar{\delta} = 1/n \sum_i \delta_i$. We know already that $E(d_i^2)$ is the same for all i, and we can thus write

$$E(d_i^2) = \frac{1}{n}\sum_k E(d_k^2) = \frac{1}{n}\sum_k E(\delta_k^2) - E(\bar{\delta}^2) \qquad \text{S30}$$

To compute $E(\bar{\delta}^2)$, we need to know $E(\delta_i\delta_j)$ and therefore $E(v_i v_j)$. To compute $E(v_i v_j)$ for $j > i$, we can write $v_j = v_i + x_{j-i}$ and understand that $x_{j-i}$ follows the same distribution as $v_i$ except that there are now only $n - i$ samples taken from an interval $[0,1 - v_i]$ and $x_{j-i}$ is the j-i[th] sample.

$$E(v_i v_j) = E\left(v_i(v_i + x_{j-i})\right) = E(v_i^2) + E\left(v_i \frac{(j-i)(1-v_i)}{n-i+1}\right) \qquad \text{S31}$$

$$= \frac{i(j+1)}{(n+1)(n+2)}$$

Using this we see that for $j > i$

$$E(\delta_i\delta_j) = E\left(\left(\frac{i}{n+1} - v_i\right)\left(\frac{j}{n+1} - v_j\right)\right) = \frac{i(n-j+1)}{(n+1)^2(n+2)} \qquad \text{S32}$$

Then we find, after a long but straightforward calculation, that

$$E(\bar{\delta}^2) = \frac{1}{n^2}\sum_i\sum_j E(\delta_i\delta_j) = \frac{1}{n^2}\sum_{i=1}^{n}\left\{\sum_{j=1}^{i-1} E(\delta_i\delta_j) + \sum_{j=i}^{n} E(\delta_i\delta_j)\right\} = \frac{1}{12n} \qquad \text{S33}$$

And therefore

$$E(d_i^2) = \frac{1}{n}\sum_k E(\delta_k^2) - E(\bar{\delta}^2) = \frac{1}{6(n+1)} - \frac{1}{12n} = \frac{n-1}{12(n+1)n} \qquad \text{S34}$$

Note that the dependence on *i* has disappeared: i.e., the variance of *d* is time-invariant; which is what we aimed to show. Also note that while we made various assumptions to simplify the above derivations, our theoretical solutions accurately predict simulated data (Figure 2-Supplement 1); showing these assumptions have little impact on the results.

If we could now compute Var[max(*d'*)], E[max(*d'*)] and E[max(*d*)] from the above solutions for *d'*, we would even be able to construct a closed-form solution for the ZETA-test's p-value in the case of exponentially-distributed inter-spike intervals. Unfortunately, the distribution of max(*d*) is unknown and fairly complex, because the elements of *d* are not statistically independent.

### Supplementary figures and legends

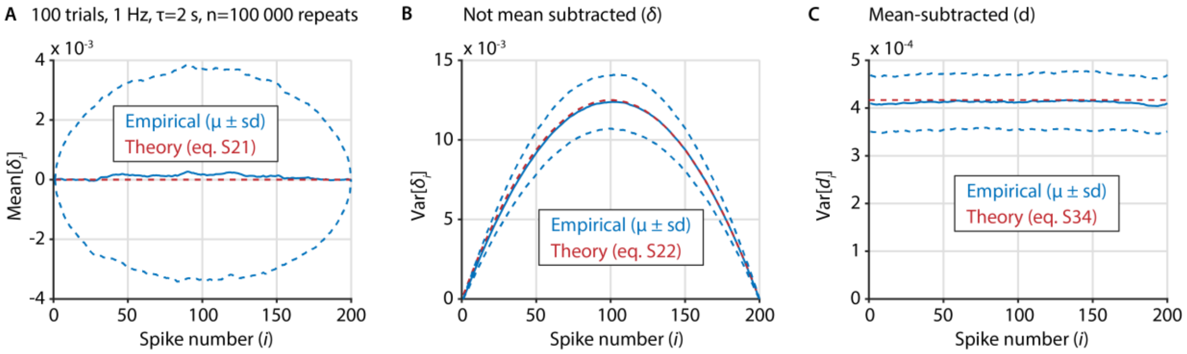


Figure 2 - Figure supplement 1. Several important variables underlying the ZETA-test can be derived analytically in the case of exponentially-distributed inter-spike intervals. A) We sampled 100 000 artificial neurons with exponentially-distributed inter-spike intervals, each spiking at 1 Hz during 100 trials that last 2 seconds. To compile data over neurons, we interpolated the spike position to [1 200], regardless of the real number of spikes. Blue shows the mean +/- standard deviation of the mean of $\delta_i$ for each spike number $i$, while the theoretical prediction is shown in red. B) Same as A, but for the variance of $\delta_i$ as a function of $i$. C) Same as B, but after recentering $\delta$ to mean-zero, showing the variance of $d_i$ as a function of $i$.

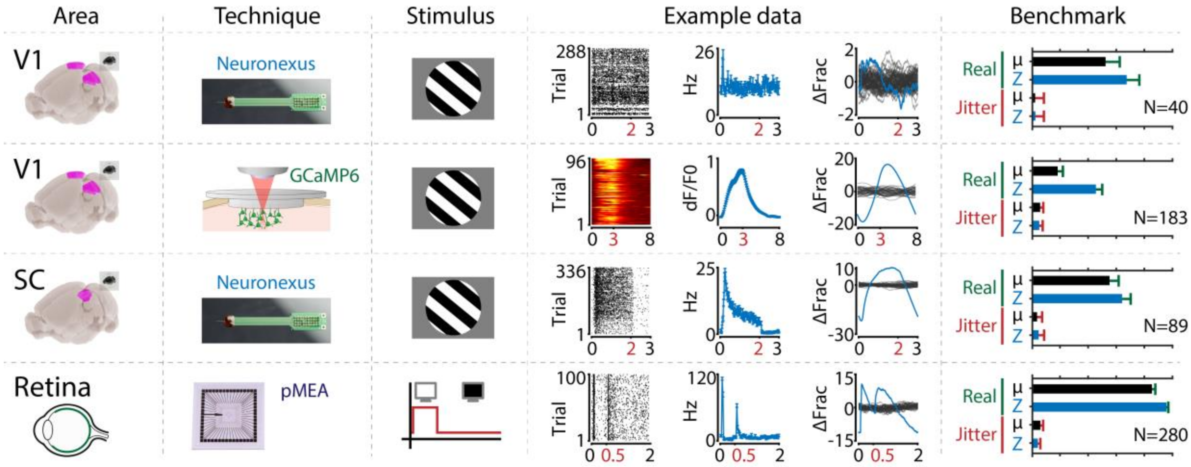


Figure 3 - Figure supplement 1. The ZETA-test is more sensitive than a mean-rate approach across a range of areas, recording techniques and stimuli. From left to right the figure shows the following. First column: the recorded area. Second column: recording technique. Third column: stimulus. Fourth column: example data including raster plot or heat map (lhs), neural activity in spiking rate (Hz) or dF/F0 (middle) and activation deviation (rhs, ΔFrac). All cells depicted here were significant using ZETA. Fifth column: benchmark performance for neuronal inclusion percentage (‘Real’, green) and false alarm rate after jittering (‘Jitter’, red) for tests using mean-rate (black, μ) and ZETA (blue, Z). From top to bottom, we investigated several data sets: mouse V1 responses to drifting gratings recorded with a Neuronexus probe (top row) and GCaMP6f (second row), mouse superior colliculus responses to drifting gratings recorded with a Neuronexus probe (third row), and zebrafish retinal ganglion cell responses to on/off light flashes recorded with a perforated micro-electrode array (fourth row). Bar graphs show inclusion rate and 75$^{th}$ percentile confidence interval.

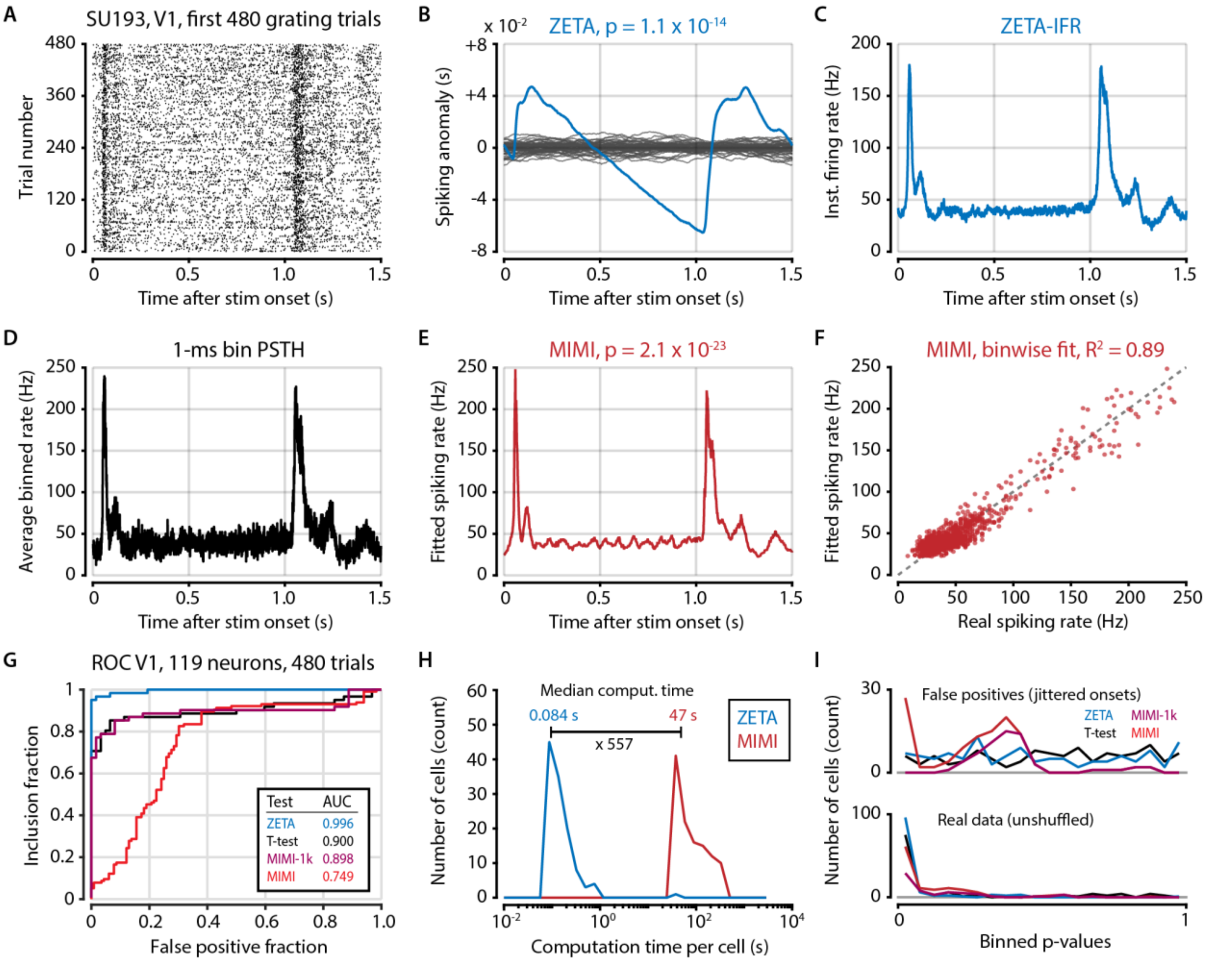


Figure 3 - Figure supplement 2. Model-based methods require high spike counts or hyperparameter tuning. A) Raster plot of example (high firing rate) V1 cell. Because of the prohibitively long computation time of fitting a multiplicative inhomogeneous Markov interval (MIMI) model to the data we opted to use only the first 480 trials recorded for each cell (N=119 neurons in V1 from 2 animals). B) Spiking anomaly (blue) and shuffle controls (grey) for the same cell, showing a clear stimulus-locked response. C) The instantaneous firing rate captures the neuron's spiking response very well. D) 1-ms binning provides a similar time resolution to both the IFR-method (panel C) and the MIMI-fit (panel E) but is much noisier at the level of single bins. E) The MIMI-model fit also captures the neuron's spiking response well (see F). Statistical significance with the MIMI-model fit was determined by calculating bin-wise *d'* values from the confidence intervals at each 1-ms bin (see Methods). F) The MIMI-model fit explains a large proportion of the variance of the PSTH ($R^2$=0.89 for this example cell), but still underperforms when used as the basis for a statistical test for stimulus-responsiveness (see panel G). G) ROC analysis showing the performance of the ZETA-test, t-test, and two versions of the MIMI-model fit. While the MIMI-model fitting worked well for cells with high spiking rates, cells with low firing rates were prone to spurious low p values during shuffle controls (see panel I). While the base MIMI-model gave an AUC of 0.749 (versus 0.996 of ZETA), excluding cells with fewer than 1000 spikes during this 480-trial epoch increased the MIMI-model's performance to on-par with the t-test (t-test AUC=0.900, MIMI-1k AUC=0.898). The threshold of 1000 spikes corresponds to an average firing rate of 1.38 Hz. H) In addition to performing worse than the ZETA-test, the MIMI-method was also much more computationally intensive: the median computation time was 557 times longer than for the ZETA-test. I) Distribution of p-values for real data (top) and shuffled controls (bottom) for the four tests; note that excluding cells with low firing rates removes all of the MIMI's false positives.

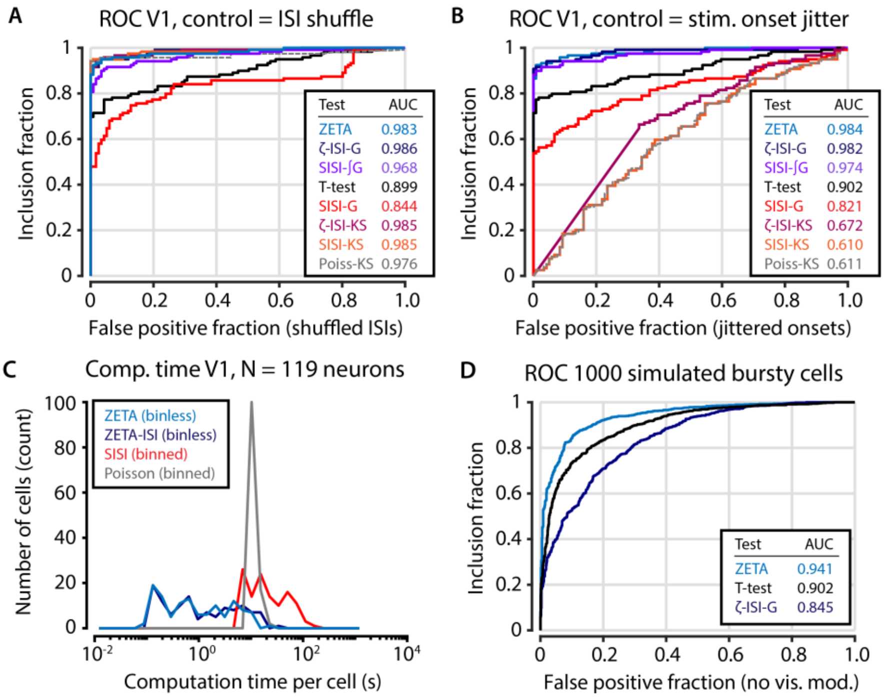


Figure 3 - Figure supplement 3. Intermediate statistical tests reveal the functions of the components of the ZETA-test. A) Various forms of statistical tests; most perform well when the test's null distribution matches the real distribution of the unresponsive population (shuffled inter-spike intervals). B) When the test's assumed (shuffled ISIs) and real (jittered onsets) null distributions are different, the Kolmogorov-Smirnov based tests fail. Of the remaining tests, the ZETA-test and ISI-shuffle ZETA-test perform best. Note that the difference in performance with Figure 3 – Figure supplement 2 is due to different data selection. C) Bin-based tests (SISI, Poisson) are slower than binless tests (ZETA, ZETA-ISI). D) Although the ZETA-test and ISI-shuffle ZETA-test performed similarly for data sets with mostly regular spiking cells (panels A,B), the ISI-shuffle ZETA-test performs considerably worse in differentiating stimulus-modulated from unmodulated bursty cells; ZETA-test AUC=0.941; Mean-rate t-test AUC=0.902; ISI-shuffle ZETA-test AUC=0.845.

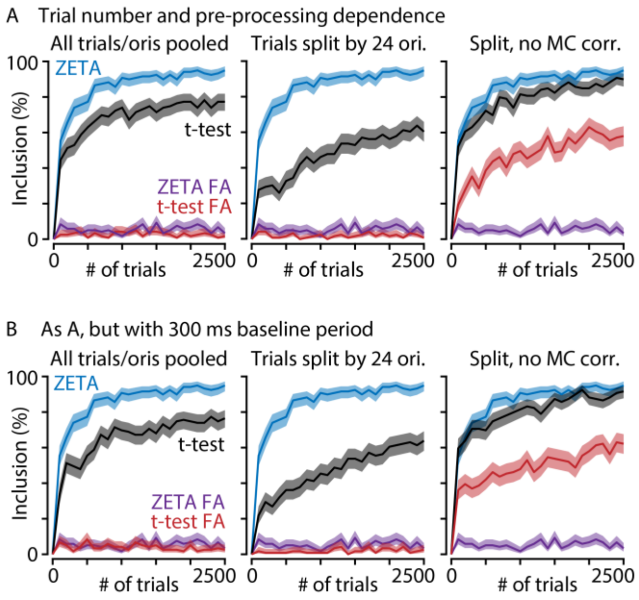


Figure 3 - Figure supplement 4. A) The ZETA-test performs better than t-tests at low trial numbers and regardless of data preparation. Graphs show V1 inclusion percentage as a function of number of trials subsampled from the whole data set, using the ZETA-test (blue) and t-test (black), and false alarms after jittering using the ZETA-test (purple) and t-test (red). Left-hand panel shows results when trials of all orientations are pooled. Middle and right-hand panels show results when t-test p-values are calculated separately per orientation (24 groups) and Bonferroni-corrected (middle) or not corrected (right). B) As panel A, but now using a 300-ms window preceding stimulus onset as baseline. There is little difference in t-test performance between using a 300-ms or 500-ms window. In all cases, the ZETA-test performs better than t-tests.

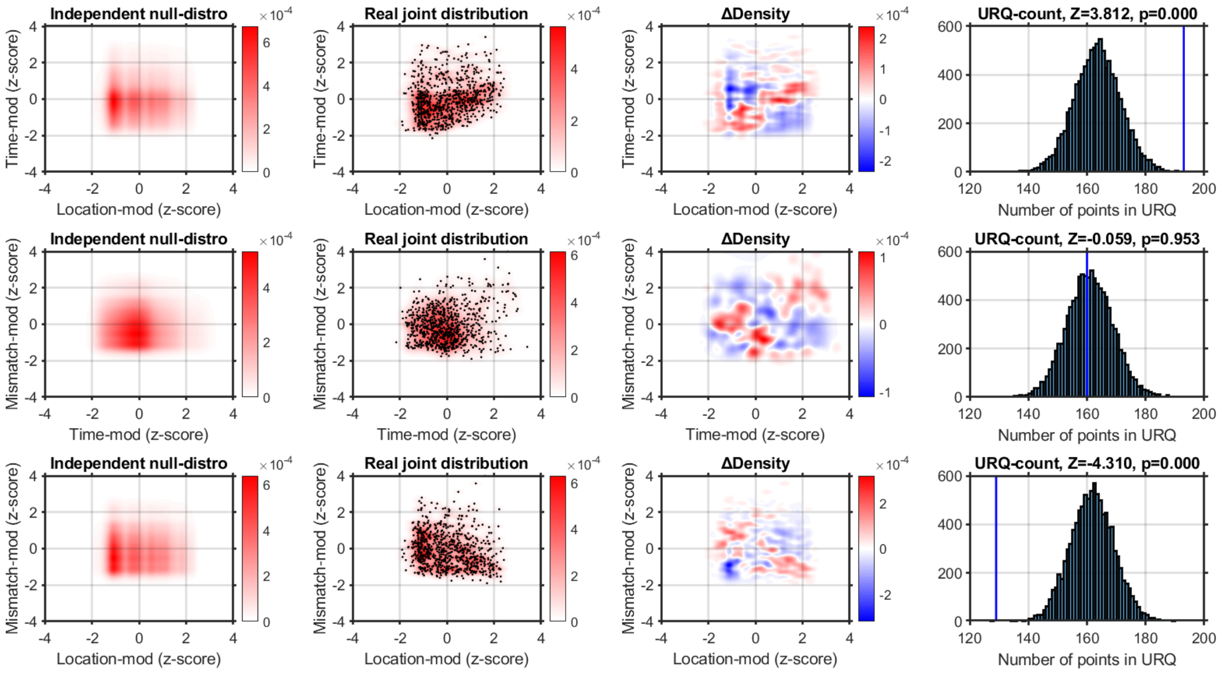


Figure 7 – Figure supplement 1. Using a kernel-density estimator (KDE) to investigate the joint-encoding of two features. Each column shows one step in our analysis procedure, and each row shows the comparison of two features. Top row shows the comparison of location-modulation (x-axis) with time-modulation (y-axis). Left-hand panel: probability density derived from combining the independent KDE-distributions of location-modulation and time-modulation. Middle-left panel: real joint probability distribution (red-scale). One point is a single neuron's modulation after z-scoring per recording. Middle-right panel: the difference in probability density between the real and null-hypothesis distributions. Blue indicates fewer neurons in the real joint-distribution than in the null-hypothesis distribution. The upper-right quadrant is mostly red, indicating the real distribution contains more neurons that encode both location and time than expected by chance. Right-hand panel: we sampled 10,000 synthetic neuronal populations from the null-hypothesis probability distribution, and calculated for each synthetic population the number of neurons that fell in the upper-right quadrant (URQ): i.e., the number of neurons that show both above-average location-modulation as well as above-average time-modulation. The histogram shows the number of neurons in the URQ for each of the 10,000 populations, and the blue vertical line shows that the real number of neurons in the URQ is much higher than expected by chance ($z=3.812$, $p=1.144 \times 10^{-4}$). Middle row: as top row, but now comparing time- and mismatch-modulation. The URQ-analysis showed that time- and mismatch-modulation were not more often seen in the same neurons than expected by chance if the two features are independently encoded ($z=-0.059$, $p=0.953$ (n.s.)). Bottom row: location on the virtual track and visuomotor mismatch are less likely to be encoded by the same neuron than if the two features were randomly distributed ($z=-4.310$, $p=2.019 \times 10^{-5}$). This shows that neurons show an encoding specialization for either spatial location or visuomotor mismatch, but not both.

Analyzing the Pearson correlation per recording we find qualitatively similar results. Mean Pearson correlation between time-modulation and location-modulation across recordings: $n=7$, mean $r=0.2272$; one-sample t-test vs. 0: $p=0.0164$. Mean correlation between time- and mismatch-modulation: $r=0.1267$, $p=0.2829$ (n.s.). Mean correlation between location- and mismatch-modulation: $r=-0.2249$, $p=0.0414$.
